# Urolithin B reduces the aggregate load of islet amyloid polypeptide in *Caenorhabditis elegans*

**DOI:** 10.1101/2025.07.01.662492

**Authors:** Mehmet Akdag, Sofia Ferreira, Regina Menezes, Tessa Sinnige

## Abstract

The progressive loss of pancreatic β-cells is one of the defining features of Type 2 Diabetes Mellitus (T2DM), and is thought to be driven by the aggregation of islet amyloid polypeptide (IAPP). This highly amyloidogenic pancreatic hormone is co-secreted with insulin, and its elevated secretion can lead to toxic fibrillar aggregation. Despite numerous studies focusing on understanding the molecular mechanisms of IAPP aggregation, few therapeutic strategies exist to counter its toxicity. Urolithin B is a natural metabolite derived from the digestion and intestinal microbiota action on ellagitannin-rich foods. This compound was suggested to counteract IAPP aggregation and toxicity *in silico* and in yeast models expressing human IAPP. In this study, we focused on the protective potential of urolithin B using our previously characterised transgenic *Caenorhabditis elegans* IAPP-GFP model. We report that urolithin B reduces the levels of insoluble IAPP-GFP in the body wall muscle cells, while the mitochondrial association of IAPP-GFP remains unaltered. We show that the *C. elegans* model exhibits a reduced lifespan compared to controls, providing *in vivo* evidence of the toxic effects associated with IAPP-GFP expression. The lifespans of IAPP-GFP versus control animals were differentially modulated by urolithin B treatment, suggesting an interplay between the metabolite and IAPP-GFP mediated toxicity. To further elucidate the mode of action of urolithin B, we conducted *in vitro* assays and found no evidence of direct interaction with lipid-associated IAPP, suggesting that its effects are not mediated by interference with IAPP–membrane interactions. These results pave the way for further therapeutic developments targeting IAPP aggregation in T2DM.

## Introduction

Over 500 million people globally suffer from diabetes [1]. The most prevalent form is Type 2 Diabetes Mellitus (T2DM), which accounts for approximately 90% of total cases of the disease. Key pathological hallmarks of T2DM include insulin resistance, hyperglycaemia, and amyloid deposition in the pancreas. Due to insulin resistance, the levels of insulin and the related hormone islet amyloid polypeptide (IAPP, also known as amylin) are elevated in T2DM.

IAPP is a peptide hormone produced by pancreatic β-cells, where it is co-expressed, co-stored, and co-secreted with insulin [2]. The major physiological functions of IAPP include gastric emptying and the regulation of glucose metabolism. It is synthesised as an 89-amino-acid-long preprohormone (preproIAPP), with the signal peptide being cleaved off upon reaching secretory organelles, producing the 67-amino-acid-long proIAPP. ProIAPP is a zymogenic prohormone that undergoes further processing prior to secretion. The flanking regions of the peptide are excised by prohormone convertases and the carboxypeptidase (CPE), the C-terminus is amidated, and finally, an intramolecular disulfide bridge is formed between the cysteine residues 2 and 7 to produce the 37-residue-long active mature IAPP hormone [3–5]. Because of its highly amyloidogenic nature, IAPP levels are strictly regulated in normal physiological conditions. However, in T2DM, increased insulin demand leads to hyperamylinemia, which is a trigger for oligomerisation and amyloid fibril formation.

The aggregation of IAPP is believed to result in pancreatic toxicity, death of β-cells, and ultimately organ failure in T2DM [6–8]. Consequently, understanding and targeting IAPP aggregation has become a focus for controlling T2DM pathology. Numerous approaches have been proposed to manipulate IAPP toxicity, including therapies aimed at improving the survival of pancreatic β-cells (such as GLP-1 receptor agonists) [9–11], small molecule inhibitors of fibril formation[12], immunotherapeutic agents [13–15], and activation of the chaperone machinery to enhance protein homeostasis [16,17].

In addition to these strategies, dietary (poly)phenols have emerged as potent inhibitors that can prevent IAPP aggregation and associated toxicity. Compounds like epigallocatechin gallate (from green tea) [18,19], resveratrol (from red grapes) [20,21], and curcumin (from turmeric) [22] have been recognised as candidates for slowing IAPP aggregation and protecting pancreatic β-cells functionality.

Given that dietary (poly)phenols suffer extensive metabolism in the human body, particularly by gut microbiota, attention has increasingly shifted toward their bioactive metabolites. These metabolites often differ significantly from their parent compounds in terms of bioavailability, stability, and biological activity. As such, targeting these downstream products offers a more physiologically relevant approach to identifying effective modulators of IAPP aggregation and toxicity, better reflecting the compounds that are present and active *in vivo* [23].

A recent *in silico* screen performed by us pointed out urolithin B [24], a metabolite derived from ellagitannin-rich foods such as pomegranates and berries [23], as a potential inhibitor of IAPP aggregation and toxicity. Cell-free studies demonstrated that urolithin B delays the kinetics of IAPP aggregation and alters the structure and morphology of the resulting aggregates and amyloid fibrils [24]. The metabolite was also shown to reduce IAPP aggregation and to mitigate IAPP-induced toxicity in a yeast model [24]. Nevertheless, the bioactivity of urolithin B against IAPP aggregation and IAPP-induced proteotoxicity remains unexplored in more complex animal models.

The nematode *Caenorhabditis elegans* is a well-characterised model organism extensively utilised to study human disease mechanisms. Its short lifespan and optically transparent nature make it an ideal *in vivo* system to investigate protein aggregation. Recently, we developed a *C. elegans* model expressing IAPP-GFP, which featured mitochondrial localisation and partial accumulation into insoluble aggregates [25]. Here we employ this model to assess the bioactivity of urolithin B on IAPP aggregation *in vivo*, complemented by *in vitro* assays.

## Results

### Urolithin B attenuates IAPP-GFP accumulation in *C. elegans* body wall muscle cells

To assess the cytotoxicity and bioactivity of urolithin B *in vivo*, we used our previously characterised *C. elegans* model expressing IAPP-GFP in the body wall muscle cells. We previously reported that the IAPP-GFP signal decorates the body wall muscle cells with elongated and spherical foci, along with a certain level of diffuse IAPP-GFP signal [25]. On the other hand, GFP-expressing control animals present a completely diffuse signal pattern.

To determine whether urolithin B treatment affects IAPP-GFP expression and localisation, synchronised animals were maintained on urolithin B-supplemented plates (100 *µ*M and 200 *µ*M) and mock-treated plates. UV-inactivated *Escherichia coli* was used as a food source to prevent urolithin B from being metabolised by the bacteria. Animals were maintained until they reached day 3 of adulthood. GFP animals showed a diffuse fluorescence signal throughout the muscle cells, and urolithin B treatment did not alter the GFP distribution even at the highest concentration (**Figure 1A**). Additionally, we did not observe any significant difference in the GFP signal intensity resulting from urolithin B treatment (**Figure 1B**), suggesting that urolithin B does not affect GFP levels or its localisation.

**Figure 1.**
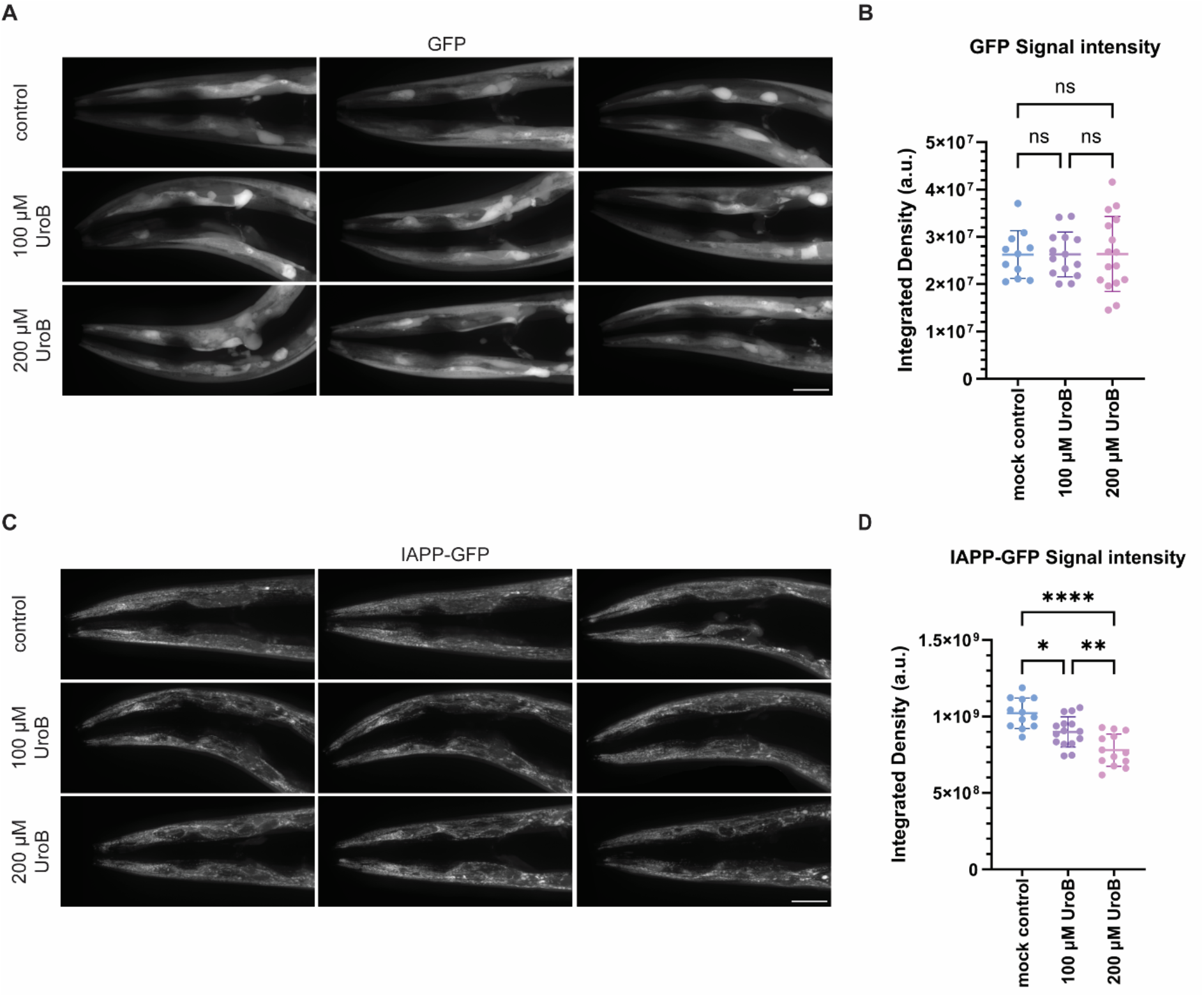
Urolithin B treatment decreases the IAPP-GFP signal in body wall muscle cells. (**A**) Representative images of GFP-expressing animals treated with increasing urolithin B (0, 100, and 200 *µ*M) concentrations. (**B**) Quantification of total GFP fluorescence in the body wall muscle cells in the head region. (**C**) Representative fluorescence images of IAPP-GFP expressing animals under the same treatment conditions. (**D**) Quantification of total IAPP-GFP signal in the body wall muscle cells in the head region. Scale bars 25 *µ*m, One-way ANOVA analysis with Tukey multiple comparison test was employed for statistical analysis (ns not significant; * p < 0.05; ** p < 0.01, ****p<0.0001). Data are presented as mean ± SD.

We then evaluated IAPP-GFP expression and localisation using microscopy. IAPP-GFP animals maintained their distinct spherical and elongated foci structures in the body wall muscle cells after urolithin B treatment (**Figure 1C**). However, the IAPP-GFP signal in the head region declined with increased urolithin B concentrations, suggesting a decrease in IAPP-GFP protein levels (**Figure 1D**).

### Urolithin B does not disturb mitochondrial localisation of IAPP-GFP

Urolithins have previously been reported to act on mitochondria by inducing mitophagy [26,27]. Our previous work demonstrated partial co-localisation of IAPP-GFP with mitochondria using a TOMM20 mitochondrial marker strain [25]. We investigated whether the decrease in IAPP-GFP level correlates with that of the mitochondria following urolithin B treatment (**Figure 2A**). The analysis revealed that the overall mitochondrial signal level in the head region of the muscle cells was not altered with increased urolithin B treatment levels (**Figure 2B**). We also quantified IAPP-GFP co-localisation with mitochondria, demonstrating that the interaction between IAPP-GFP and mitochondria remained unaffected (**Figure 2C**). These results suggest that urolithin B treatment causes a concentration-dependent decrease in overall IAPP-GFP signal, without affecting its co-localisation to mitochondria or mitochondrial levels.

**Figure 2.**
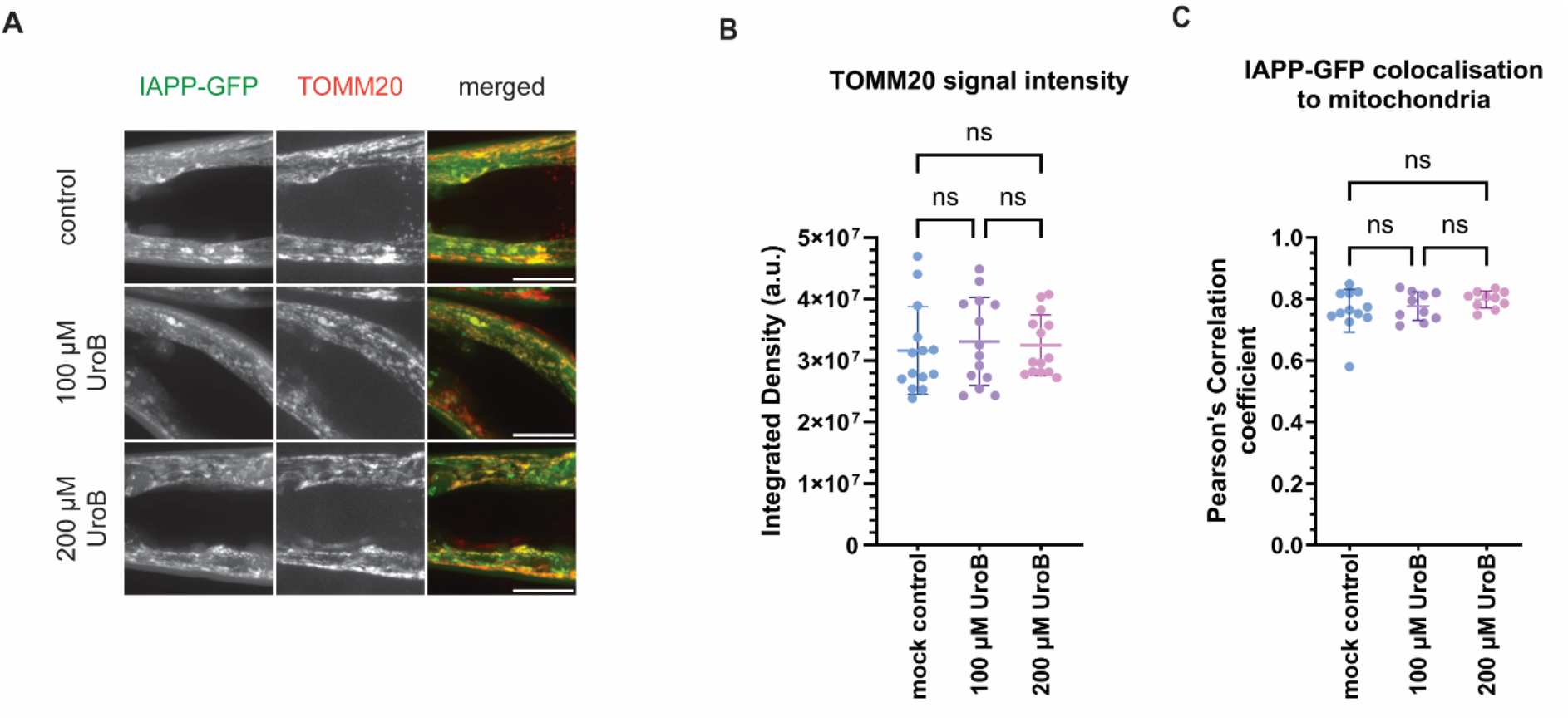
Urolithin B treatment does not alter IAPP-GFP mitochondrial interaction in the body wall muscle cells. (**A**) Representative images of IAPP-GFP (green) and mitochondrial marker TOMM20-mKate (red) expressing animals treated with or without urolithin B. (**B**) Quantification of total TOMM20-mKate fluorescence signal in the head region of the body wall muscle cells treated as in A. (**C**) Pearson’s correlation analysis of IAPP-GFP co-localisation to the mitochondria. Scale bars 25 *µ*m, One-way ANOVA analysis with Tukey multiple comparison test was employed for statistical analysis (ns not significant). Data are presented as mean ± SD.

### Urolithin B reduces insoluble IAPP-GFP load in *C. elegans*

We proceeded with biochemical fractionation experiments to investigate whether urolithin B is capable of lowering insoluble IAPP-GFP aggregate levels in *C. elegans*, as observed in yeast models [24]. We treated age-synchronised animals with urolithin B until day three of adulthood and fractionated the lysates into detergent-soluble and -insoluble fractions (**Figure 3A**). As we previously showed in yeast [24] and *C. elegans* [25], IAPP-GFP partially accumulates in the insoluble fraction in the mock-treated control group. However, we noted a dose-dependent reduction in the urolithin B-treated IAPP-GFP insoluble fraction (**Figure 3B**). This result suggests that urolithin B acts to reduce the IAPP-GFP aggregate load.

**Figure 3.**
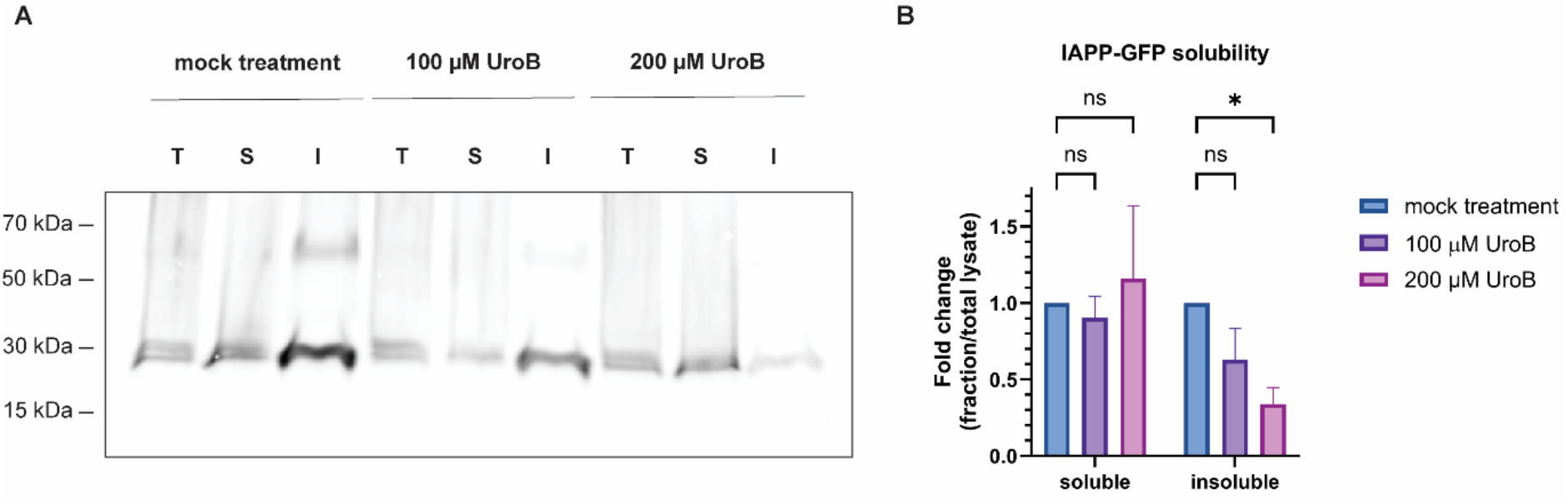
Urolithin B treatment reduces insoluble IAPP-GFP aggregates. (**A**) Representative western blot showing detergent-soluble and insoluble fractions of IAPP-GFP lysates probed with anti-GFP antibody. Lysates were prepared from IAPP-GFP animals kept on 0, 100, or 200 *µ*M urolithin B-containing plates until day three of adulthood. Total lysate (T), soluble fraction (S), and insoluble fraction (I) are shown. (**B**) Quantification of fractionation assays. Detergent-soluble and -insoluble fractions are normalised to the total lysate. Data are visualised as fold change compared to the mock-treated group. The experiment was done in triplicate, and the data are represented as mean ± SD (ns not significant; * p < 0.05).

### Urolithin B differentially affects lifespan in IAPP-GFP and GFP control worms

We next investigated whether the urolithin B-mediated reduction in insoluble IAPP-GFP influences the health of the *C. elegans* models. We performed lifespan assays in both IAPP-GFP and GFP control animals and found that IAPP-GFP animals have a shortened lifespan compared to the control, the median lifespans being 14 and 19 days, respectively (**Figure 4A**). These results highlight the toxicity associated with IAPP-GFP accumulation. Unexpectedly, urolithin B treatment caused a reduction in the average lifespan of GFP control animals from 19 days to 15.5 days at 100 *µ*M (**Figure 4B**). The reduction was not concentration-dependent, suggesting that the toxicity had already reached a plateau at 100 *µ*M (**Figure S1**). However, these data indicate a certain level of toxicity associated with urolithin B treatment of healthy control animals. A similar effect was observed for wild-type *C. elegans*, indicating that the effect is not related to GFP expression (**Figure S1**).

**Figure 4.**
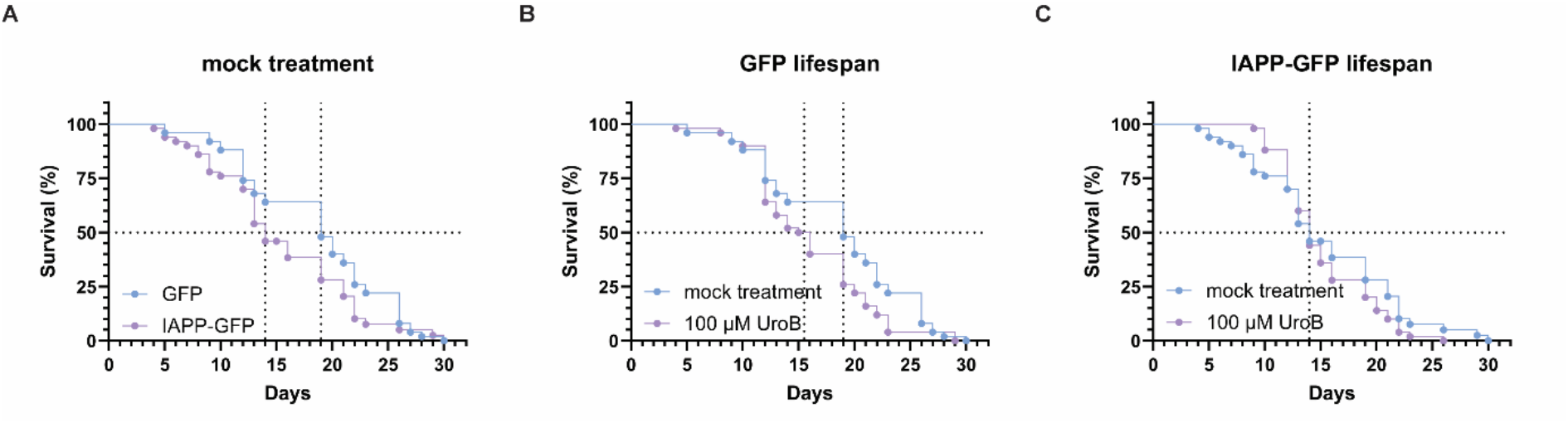
Urolithin B differentially affects the lifespan of IAPP-GFP animals compared to GFP control. (**A**) Lifespan of GFP control and IAPP-GFP expressing animals without urolithin B treatment. (**B**) The effect of urolithin B treatment (100 *µ*M) on GFP-expressing nematodes. (**C**) The effect of urolithin B treatment (100 *µ*M) on IAPP-GFP animals’ lifespan. 50 animals were scored for each condition. Dashed lines represent the median lifespan of the populations. The experiment was performed in a blinded manner, and the lifespans were determined using Kaplan-Meier analysis.

By contrast, we did not observe a reduction in lifespan in IAPP-GFP animals treated with urolithin B at 100 *µ*M (**Figure 4C**) or 200 *µ*M (**Figure S1C**). Urolithin B-treated IAPP-GFP animals performed comparably to mock-treated animals, with a median lifespan of 14 days (**Figure 4C**). While urolithin B does not restore the lifespan deficit caused by IAPP-GFP, its beneficial role in reducing the IAPP-GFP aggregate load may be masked by the toxic effect that is also observed in the control animals. As such, the differential effect of urolithin B on IAPP-GFP versus GFP lifespan is suggestive of an interplay between the compound and IAPP-GFP mediated toxicity.

### Urolithin B does not alter IAPP-mediated membrane disruption *in vitro*

Prompted by our findings indicating an association between urolithin B bioactivity and IAPP, we further explored the possible mechanism using *in vitro* approaches. The recruitment and fibrillation of IAPP on membranes are proposed to be one of the toxicity mechanisms of IAPP. To investigate the effect of urolithin B on the IAPP-lipid interaction, we prepared large unilamellar vesicles (LUVs) composed of DOPC and DOPS lipids (7:3 molar ratio) and measured the aggregation behaviour of IAPP in the presence of lipids and urolithin B (**Figure 5A**). However, we did not observe a urolithin B-specific change in IAPP aggregation kinetics at a 1:100 peptide-to-lipid molar ratio.

**Figure 5.**
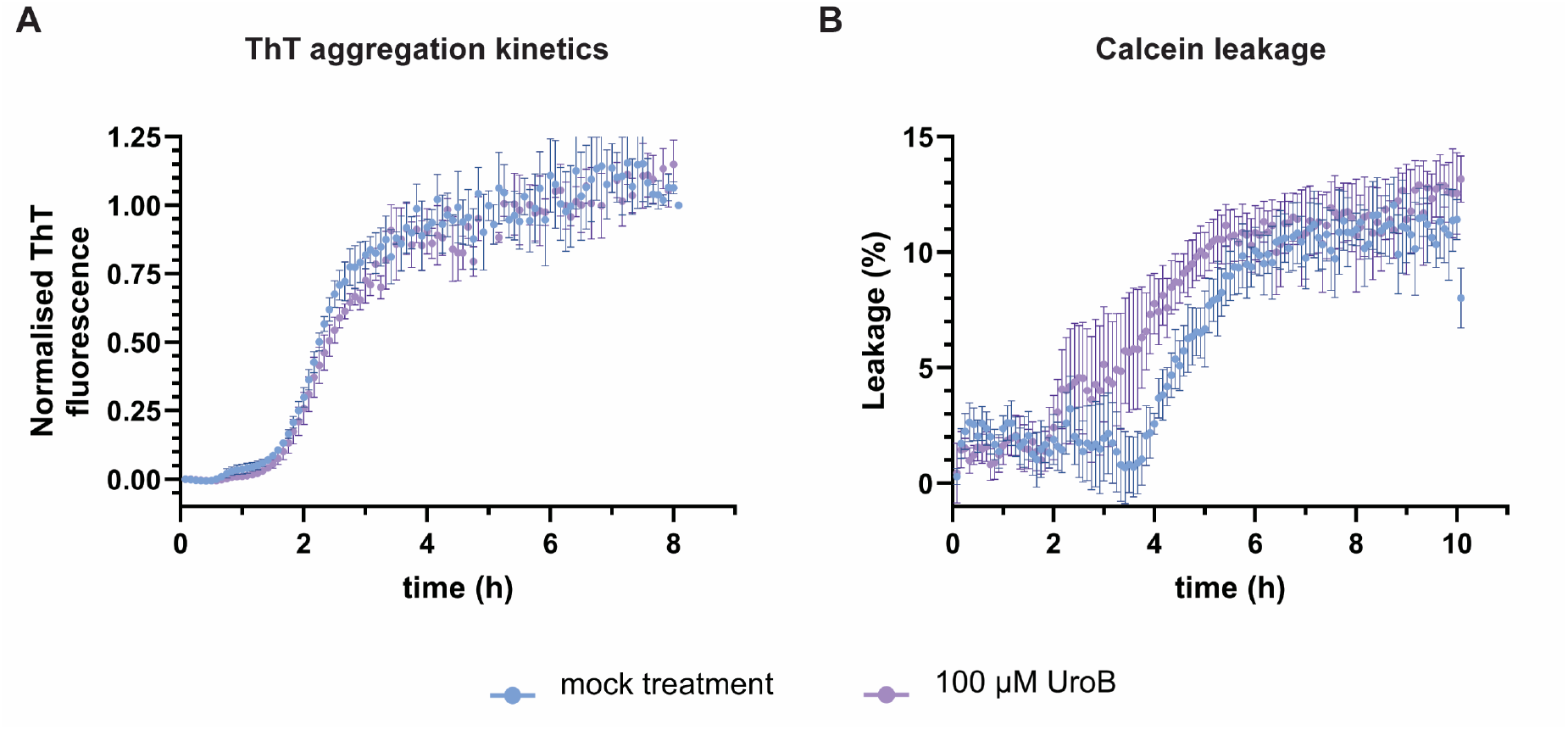
Urolithin B treatment does not affect the interaction between IAPP and model membranes *in vitro*. (**A**) ThT assay showing IAPP aggregation kinetics in the presence of DOPC:DOPS (7:3) lipid vesicles, with and without the addition of 100 *µ*M urolithin B. (**B**) Calcein leakage assay showing the membrane permeabilisation kinetics of IAPP in the absence and presence of urolithin B. 5 *µ*M IAPP, 100 *µ*M urolithin B, and 500 *µ*M LUVs (DOPC: DOPS 7:3 ratio) were used, and each assay was performed in triplicate. The percentage of calcein leakage was normalised to the fluorescence of Triton X-100 leakage. The data are represented as mean ± SEM.

Next, we employed a calcein leakage assay to evaluate the effect of IAPP on membrane integrity upon urolithin B treatment. IAPP is known to induce leakage in model membranes, with kinetics corresponding to that of fibril formation [28,29]. Although we observed a slight shift between the two curves, the kinetics of leakage do not appear to be delayed by urolithin B treatment, and the plateau value is not reduced (**Figure 5B**). Thus, urolithin B does not seem to interfere with the disruption of model membranes caused by IAPP aggregation *in vitro* under the conditions tested.

## Discussion

In this study, we examined the effect of urolithin B on IAPP aggregation using our previously characterised IAPP-GFP-expressing *C. elegans* model, along with *in vitro* assays. Urolithin B treatment led to a reduction in the total IAPP-GFP fluorescent signal in the body wall muscle cells in the head of the nematodes and caused a decrease in the amount of insoluble IAPP-GFP as seen by fractionation experiments.

We did not observe any change in mitochondrial abundance or the fraction of IAPP-GFP co-localising with mitochondria following urolithin B treatment. These results suggest that the reduction in IAPP-GFP is not directly associated with mitophagy. Previous studies reported that the closely related metabolite urolithin A, and urolithin A-containing fermented microbial extracts, induced mitophagy in *C. elegans* [26,30]. Urolithin A and B differ from each other by one alcohol group, and it is not known whether urolithin B also induces mitophagy. However, several autophagy genes were previously shown to be upregulated upon urolithin B treatment in the IAPP yeast model [24]. Considering that autophagy is likely to play a role in the clearance of insoluble aggregates, its contribution to protein turnover mechanisms should be studied in more detail to further understand the link between urolithin B and IAPP-GFP insoluble aggregate clearance in the *C. elegans* model.

Unexpectedly, we found that urolithin B was toxic to healthy *C. elegans*, as seen from the shortened lifespan of the control animals. This result contrasts with the previously reported lifespan extension by several urolithins, including urolithin B [26]. Differences in experimental conditions are likely to account for this result. In the previous report, the urolithins were added to the NGM, whereas we supplemented it with the bacterial food source. As such, the effective uptake is expected to be much larger in our experiments, reaching a dose that is apparently toxic to healthy *C. elegans*. Furthermore, it was shown that UV-killing of the bacterial food source reduced the lifespan extension by urolithin A [26]. In our study, we consistently used UV-killed bacteria to avoid further metabolic conversions.

We are convinced that urolithin B toxicity is not an issue for humans, given that it is a naturally occurring metabolite. However, the dose may be a limiting factor, particularly considering its moderate bioavailability. Previous studies report that urolithin B reaches plasma concentrations of approximately 0.01 *µ*Mol/L six hours after ingestion of pomegranate juice concentrate (387 mg/L of anthocyanins, 1561 mg/L of punicalagins, 121 mg/L of ellagic acid, and 417 mg/L of other hydrolysable tannins) [31]. Although its distribution in human tissues remains unknown, animal studies have detected both aglycone and sulphate forms of urolithin B in the pancreas, liver, and heart [32]. These findings suggest a potential for peripheral bioactivity, but the lack of human data represents a significant knowledge gap that warrants further investigation.

We furthermore show that our previously established IAPP-GFP *C. elegans* model exhibits a reduced lifespan compared to controls, providing *in vivo* evidence of the toxic effects associated with IAPP-GFP expression. The survival of IAPP-GFP animals was unaffected by urolithin B treatment, hinting at a potential dual effect of the compound. A possible explanation could be that urolithin B contributes to relieving IAPP-GFP toxicity by reducing the aggregate load; however, at the concentrations tested, the compound’s toxicity may offset this beneficial effect.

Based on the *in vitro* results, urolithin B does not seem to directly interfere with the interaction between IAPP and lipid membranes. These results are in line with the association between IAPP-GFP and mitochondria that we observed *in vivo*, which was also unchanged by urolithin B. Thus, it is more likely that urolithin B acts on the cytosolic fraction of IAPP, preventing its aggregation as previously suggested [24]. In addition, urolithin B may stimulate autophagic clearance of aggregated IAPP as discussed above, altogether resulting in less aggregate accumulation.

Evidently, *C. elegans* presents certain physiological limitations as a model for studying T2DM. Nematodes do not have a homologous hormone to IAPP, and they lack a pancreas-like specialised organ responsible for hormone production. Although a *C. elegans* model can provide meaningful insights, other models, such as human β-cell cultures, would be useful to shed more light on the mechanisms by which urolithin B alleviates IAPP toxicity.

In conclusion, this study shows that urolithin B reduces the levels of insoluble IAPP-GFP aggregates in *C. elegans*. The results suggest that urolithin B does not interfere with the IAPP-membrane interaction, but acts to promote its solubility. Further studies and additional model systems are required to explore the precise interplay, paving the way for the development of metabolite-derived compounds as potent drugs for T2DM.

## Methods

### *C. elegans* strains and maintenance

Nematodes were routinely maintained on nematode growth media (NGM) plates at 20 °C, seeded with *Escherichia coli* OP50. Age synchronisation was achieved by incubating gravid adults on NGM plates at 20 °C for 2 h. Eggs were incubated at 20 °C until they reached adulthood three days after, which is defined as day 1 in this study.

The following *C. elegans* strains were used in this study:

N2 (Bristol)

TSW53 mbbIs9[*unc-54*p::GFP::*unc-54* 3’ UTR]

TSW55 mbbIs11[*unc-54*p::IAPP::GFP::*unc-54* 3’ UTR]

TSW80 mbbIs9[*unc-54*p::GFP::*unc-54* 3’ UTR], foxSi16[*myo-3*p::tomm-20::mKate2::HA::*tbb-2* 3’ UTR]

TSW81 mbbIs9[*unc-54*p::IAPP::GFP::*unc-54* 3’ UTR], foxSi16[*myo-3*p::tomm-20::mKate2::HA::*tbb-2* 3’ UTR]

SJZ47 foxSi16[*myo-3*p::tomm-20::mKate2::*tbb-2* 3’ UTR]

### Drug treatment

Urolithin B was acquired from Toronto Research Chemicals. For urolithin B treatment experiments, a 100 mM urolithin B stock solution was prepared by dissolving the compound in dimethyl sulfoxide (DMSO). To prevent bacteria from metabolising the compound, the bacterial culture was treated with a 30 mJ/cm^2^ UV light using a UVP Crosslinker CL-3000 (Analytik Jena) for 10 min. The main stock was diluted in the UV-inactivated *E. coli* OP50 bacteria culture to obtain 0 *µ*M (DMSO, mock control), 100 *µ*M and 200 *µ*M urolithin B supplemented cultures. Fresh NGM plates were seeded with these cultures as a food source.

Gravid animals were kept on these plates to lay eggs for 2 h, and their offspring were maintained on drug-containing or mock plates throughout the experiments.

### Lifespan assay

For the lifespan assay, age-synchronised animals were maintained on NGM plates with UV-inactivated bacterial lawns containing either DMSO or urolithin B. Animals were transferred to fresh plates every other day after adulthood day 1. For each condition, animals were scored as alive or dead when they were not responsive to body touches. The experiment lasted until all the animals were dead. The number of animals per condition at the start of the experiment was either 50 or 100, as indicated in the figure legends. The median lifespans were determined using Kaplan-Meier analysis.

### Fractionation

Age-synchronised nematodes (ca. 5000 animals. per condition) were collected in M9 buffer (22 mM KH_2_PO_4_, 42 mM Na_2_HPO_4_, 8.5 mM NaCl, 18.7 mM NH_4_Cl,1 mM MgSO_4_) on the second day of adulthood and snap-frozen in liquid nitrogen. Pellets were resuspended in a protease inhibitor cocktail (Roche)-containing RIPA Lysis Buffer (Thermo Fischer Scientific) and lysed by using the Tissue Lyser II (Qiagen) for 5 min at a frequency of 30 s^-1^. The sample was centrifuged at 1,000 *g* for 5 min at 4 °C to remove debris. Total protein concentrations of samples were measured by BCA assay (Thermo Fischer Scientific) and corrected to equal concentrations by dilution with lysis buffer. The samples were fractionated into supernatant and pellet fractions by centrifugation at 20,800 *g* for 30 min at 4 °C. The pellet fraction was washed with lysis buffer and then dissolved in the urea buffer (8 M urea, 2 % SDS, 50 mM DTT, 50 mM Tris-HCl pH 8.0).

The fractions were boiled in 5X sample buffer (5 % SDS, 50 % glycerol, 0.1 % Bromophenol blue, 250 mM Tris-Cl pH 6.8, and 5 % β-mercaptoethanol) at 90 °C for 10 min. Fractions were run on an SDS-PAGE gel, and proteins were transferred to PVDF membrane by using the Power Blotter System (Invitrogen). Membranes were blotted using mouse anti-GFP (JL8, Takara) and goat anti-mouse (A32730, Invitrogen) antibodies. Detection was done using an Odyssey DLx Imager (Li-Cor).

### Microscopy

Age-synchronised day 2 nematodes were immobilised on microscopy slides containing 10 mM tetramisole hydrochloride (Sigma Aldrich) on 2.5 % agarose pads. Animals were imaged within 1 h after being treated with the anaesthetic agent. Imaging was performed on a Nikon Eclipse Ti2 confocal microscope equipped with an X-Light V3 spinning disk (Crest Optics) and Nikon Plan Apo 60×A/1.40 oil DIC H oil objective. Excitation and emission wavelengths were set to 470 nm and 525/50 nm for GFP, and 555 nm and 605/70 nm for mKate imaging.

### IAPP preparation

Synthetic IAPP was obtained from Pepmic. The peptide was dissolved in hexafluoro-isopropanol (HFIP) to a concentration of 1 mM and incubated overnight at room temperature. The solution was aliquoted and then dried under nitrogen gas flow. The remaining HFIP was removed by applying high vacuum for 1 h. Aliquots were stored at -80 °C until use. Peptides were reconstituted in milliQ water prior to use to a concentration of 160 *µ*M for kinetics assays and 200 *µ*M for vesicle leakage assays.

### Lipid vesicle preparation

1,2-Dioleoyl-*sn*-glycero-3-phosphocholine (DOPC, Avanti #4235-95-4) and 1,2-dioleoyl-*sn*-glycero-3-phospho-L-serine (DOPS, Avanti #840035) lipids were used in 7:3 (DOPC: DOPS) molar ratio. To prepare this lipid ratio, DOPC and DOPS stocks were prepared in methanol: chloroform (1:2 volume ratio) buffer. After measuring lipid concentrations by the Rouser assay[33], appropriate amounts of lipids were mixed.

Lipids were dried under nitrogen gas flow, and the remaining solvents were removed by incubation in a 42 °C water bath. Lipids were resuspended in Tris buffer (10 mM Tris-HCl pH 7.4, 100 mM NaCl), leading to the formation of multilamellar vesicles (MLVs). MLVs were incubated at room temperature for 1 h and resuspended once every 10 min. MLVs were subsequently homogenised by 10 freeze-thaw cycles in dry ice-cold ethanol, followed by lukewarm water, vortexing after each cycle. The MLV mixture was extruded through a 200 nm pore filter (Anotop 10, Whatman) 21 times by using an Avanti Lipid extruder kit to obtain large unilamellar vesicles (LUVs). The lipid concentration of LUVs was measured using the Rouser assay.

### Thioflavin T Assay

Thioflavin T (ThT) assays were carried out in triplicates in final volumes of 200 µL. The reaction mixtures contained IAPP (5 *µ*M), urolithin B (100 *µ*M), LUVs (500 *µ*M), Tris buffer (10 mM Tris-HCl pH 7.4, 100 mM NaCl) and ThT (20 *µ*M). A 96-well flat bottom plate (CELLSTAR black, Greiner) covered with a transparent cover sticker (Viewseal selar, Greiner) was used for the experiment. The ThT assay was performed in a climate controled room at 20 °C using a Clariostar plus (BMG Labtech) plate reader. The assay was started after an initial shaking step for 20 s at 500 rpm. 440/15 nm and 485/20 nm wavelengths were used for excitation and emission, respectively.

### Calcein leakage assay

Vesicles were prepared as stated above with the following modifications for the calcein leakage assay. MLV formation was performed by resuspending lipids in calcein solution (100 mM calcein in ultrapure water pH 7.4). The calcein stock solution was passed through a 200 nm cellular acetate membrane and the osmolality of the calcein buffer was measured by using a K-7400S Semi-Micro Osmometer (KNAUER). Osmolality was set to 250 (± < 5 %) mOsm/kg by diluting the buffer. LUVs were prepared as described above and excess dye was removed by using a 10 mL gravity size exclusion column which was equilibrated in Tris buffer (10 mM Tris-HCl pH 7.4, 118 mM NaCl) with a similar osmolality. Lipid concentration was measured using the Rouser assay.

For the leakage assay, Tris buffer, calcein vesicles, urolithin B, and IAPP peptide were used at a volume ratio of 135:50:10:5, respectively. The final concentrations were 500 *µ*M for lipids, 5 *µ*M for IAPP and 100 *µ*M for urolithin B. IAPP was substituted by ultrapure water as mock treatment and by 10 % Triton X-100 as a positive control. The excitation and emission wavelengths were set to 475/10 nm and 520/10 nm, respectively. The assay was performed in triplicate with a total volume of 200 µL using a Clariostar plus (BMG Labtech) plate reader in a climate-controlled room at 20 °C. The results were normalised by setting the Triton X-100 treatment to 100 % leakage.

### Statistical analysis

For statistical analysis, GraphPad Prism 10 was used. One-way ANOVA analysis with Tukey test was employed for microscopy analysis. Two-way ANOVA analysis with Tukey test was employed for analysing western blot results. Kaplan-Meier analysis was used for the lifespan analysis. Data are presented as the mean ± standard deviation. p ≤ 0.05 was used as the threshold for considering the data statistically significant.

## Acknowledgements

We thank Martin Haase at Physical and Colloidal Chemistry (Utrecht University) for the use of the confocal microscope. We thank Antoinette Killian, Hugo van Ingen and Sinnige lab members for valuable discussions. This work was supported by a start up grant funded by Utrecht University to T.S. This research was also funded by national funds through FCT—Foundation for Science and Technology, I.P. (Portugal), under the [DOI 10.54499/UIDB/04567/2020] and [DOI 10.54499/UIDP/04567/2020] projects. R.M. is funded by a FCT Scientific Employment Stimulus contract [DOI: 10.54499/CEECINST/00002/2021/CP2788/CT0004]. S.F. is funded by the FCT PhD grant UI/BD/151421/2021 and received a Short-Term Scientific Mission grant funded by the COST Action CA20113.

## Author contributions

M.A. and S.F. performed experiments and drafted the manuscript. All authors analysed data and reviewed and edited the manuscript.

## Data availability

All data associated with this study are available in the manuscript and supplementary information.

