## Supplementary Figure 1 for "Urolithin B reduces the aggregate load of islet amyloid polypeptide in *Caenorhabditis elegans*"

1- Molecular Biophysics and Biochemistry, Bijvoet Centre for Biomolecular Research, Utrecht University, Padualaan 8, 3584 CH Utrecht, the Netherlands
2- CBIOS, ECTS, Lusófona University, Lisbon, Portugal
3- Universidad de Alcalá, Escuela de Doctorado, Departamento de Ciencias Biomédicas, Madrid, Spain

* equal contribution

### co-corresponding authors

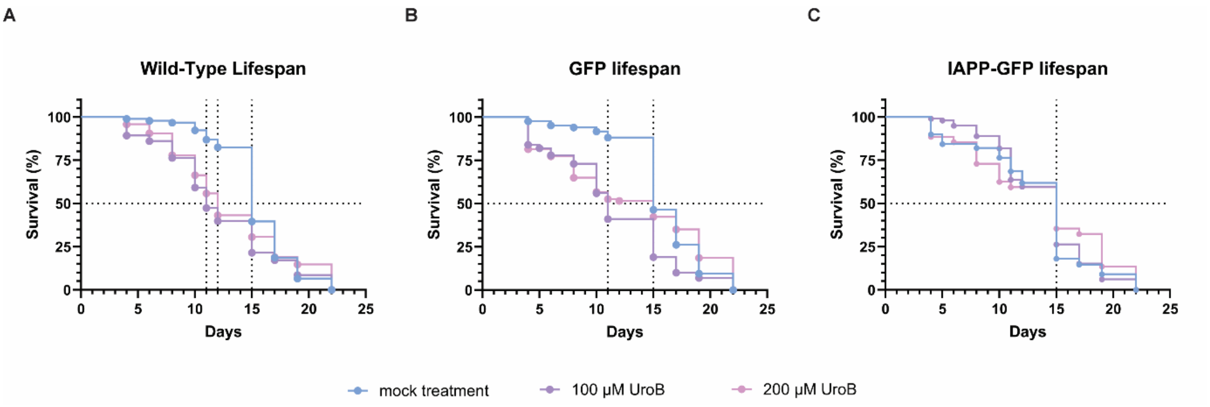


**Figure S1 The effects of urolithin B on *C. elegans* lifespan do not change beyond 100 $\mu$M.** (**A**) The effect of different urolithin B concentrations (0, 100 or 200 $\mu$M) on the lifespan of wild-type animals. (**B**) The effect of different urolithin B concentrations (0, 100 or 200 $\mu$M) on GFP-expressing nematodes. (**C**) The effect of different urolithin B concentrations (0, 100 or 200 $\mu$M) on IAPP-GFP-expressing nematodes. One hundred animals were scored for each condition. Age-synchronised wild-type animals are treated with 0 $\mu$M (blue), 100 $\mu$M (purple), or 200 $\mu$M (pink) urolithin B. Dashed lines represent the median lifespan of the populations. The experiment was performed in a blinded manner, and the lifespans were determined using Kaplan-Meier analysis.
